# A Fluorescence-Based, T5 Exonuclease-Amplified DNA Cleavage Assay for Discovering Bacterial DNA Gyrase Poisons

**DOI:** 10.1101/2023.10.16.562555

**Authors:** Matthew Dias, Trisha Chapagain, Fenfei Leng

## Abstract

Fluoroquinolones (FQs) are potent antibiotics of clinical significance, known for their unique mechanism of action as gyrase poisons, which stabilize gyrase-DNA cleavage complexes and convert gyrase into a DNA-damaging machinery. Unfortunately, FQ resistance has emerged, and these antibiotics can cause severe side effects. Therefore, discovering novel gyrase poisons with different chemical scaffolds is essential. The challenge lies in efficiently identifying them from compound libraries containing thousands or millions of drug-like compounds, as high-throughput screening (HTS) assays are currently unavailable. Here we report a novel fluorescence-based, T5 exonuclease-amplified DNA cleavage assay for gyrase poison discovery. This assay capitalizes on recent findings showing that multiple gyrase molecules can simultaneously bind to a plasmid DNA molecule, forming multiple gyrase-DNA cleavage complexes on the same plasmid. These gyrase-DNA cleavage complexes, stabilized by a gyrase poison, can be captured using sarkosyl. Proteinase K digestion results in producing small DNA fragments. T5 exonuclease, selectively digesting linear and nicked DNA, can fully digest the fragmented linear DNA molecules and, thus, “amplify” the decrease in fluorescence signal of the DNA cleavage products after SYBR Green staining. This fluorescence-based, T5 exonuclease-amplified DNA cleavage HTS assay is validated using a 50-compound library, making it suitable for screening large compound libraries.

## Introduction

Bacterial DNA gyrase, a type IIA DNA topoisomerase, is an essential enzyme for bacteria (1-6). It contains two distinct subunits, GyrA and GyrB that form an active tetrameric A_2_B_2_ complex (2,7). GyrA carries an active tyrosine residue used in DNA cleavage and religation reactions for DNA supercoiling activity (2). GyrB contains an ATP-binding site that is also required for DNA supercoiling (3). The absence of DNA gyrase in human cells makes this enzyme a highly promising and valuable target for the discovery and development of novel antibiotics (8-10). Presently, there are two main classes of gyrase inhibitors: catalytic and poisons (10,11). Catalytic inhibitors, such as novobiocin, primarily target the ATP binding site of GyrB, inhibiting the gyrase’s supercoiling activity (12-14). Unfortunately, novobiocin was withdrawn from the US market due to an unfavorable efficacy and safety profile (12). Several pharmaceutical companies have also synthesized a vast array of compounds with anti-GyrB ATPase and antibacterial activities over the past three decades (12). However, no synthetic GyrB inhibitors has been successfully developed into antibiotics yet (12).

Gyrase poisons, such as fluoroquinolones (FQs), target the active tyrosine site of GyrA through stabilizing the gyrase-DNA cleavage-complex intermediates during DNA supercoiling cycle (15-19). This gyrase poisoning mechanism makes FQs among the most effective antibiotics (20-23). Unfortunately, bacterial resistance to FQs has emerged (15,24-31) and makes the development of new, more effective antibiotics a great urgency (26). Additionally, FQs have been explored extensively (10,32,33). The limits and potential of FQs likely have been reached (10,32,33). Furthermore, the use of FQs is associated with serious side effects (34), including tendonitis and tendon rupture (35-37), peripheral neuropathy (38), hyperglycemia (39), and aortic complications (40-42). Consequently, FDA has issued multiple warnings for the use of FQs and implemented black box warnings on all FQs (43,44). In light of these challenges, there is an evident need to discover novel types of compounds targeting bacterial DNA gyrases to effectively treat bacterial infections. Examples of this ongoing effort include novel bacterial topoisomerase inhibitors (NTBIs), such as gepotidacin and zoliflodacin (45,46). These NTBIs, currently under phase 3 clinical trials for the treatment of gonorrhea, also poison gyrase, albeit targeting a different site on gyrase-DNA complexes (46,47). These new gyrase poisons offer great promise to fight “superbugs” that are resistant to almost all antibiotics (48) and avoid facing a future pandemic of untreatable bacterial infections (49).

An effective approach for discovering new DNA gyrase poisons is through high-throughput screening (HTS) of compound libraries that contain thousands or millions of drug-like compounds. However, there are currently no HTS assays that can identify or discover DNA gyrase poisons. Instead, labor-intensive and time-consuming agarose or PAGE gel-based assays are typically used to identify or confirm gyrase poisons (50,51). Nevertheless, several HTS assays are available to identify gyrase inhibitors that target the DNA supercoiling activity of gyrase (50,52-55). For example, we recently established a miniaturized, automated ultra-high-throughput screening (uHTS) assay based on the supercoiling-dependent fluorescence quenching (SDFQ) assay of DNA topoisomerases (56). We screened the NIH’s Molecular Libraries Small Molecule Repository (MLSMR) library, which contains 370,620 compounds, and identified ∼3,000 DNA gyrase inhibitors (50). Although most of these newly identified gyrase inhibitors are catalytic inhibitors, several are poisons. This was confirmed by performing agarose-gel based DNA cleavage assays by gyrase (50). We also pioneered another HTS assay, a T5 exonuclease (T5E) AT-hairpin-based HTS assay, to identify gyrase inhibitors (52). This assay is based on a unique property of T5E that can completely digest supercoiled plasmid pAB1 containing an “AT” hairpin structure and spare relaxed pAB1 (52). This HTS assay can also be converted into a miniaturized, automated uHTS assay for DNA topoisomerases.

In addition to bacterial DNA gyrase, other DNA topoisomerases are also important drug targets since DNA topoisomerases are essential enzymes (1-3,8). Currently, all clinically relevant drugs targeting DNA topoisomerases act as poisons (8,57). For example, doxorubicin and etoposide are poisons to human DNA topoisomerase IIα (8). Camptothecin and analogs kill cancer cells through poisoning human DNA topoisomerase I (8). As mentioned above, FQs, such as ciprofloxacin, are poisons against bacterial DNA gyrase and topoisomerase IV (8,15,21). A recent study demonstrated the effectiveness of cyanotriazoles as DNA topoisomerase II poisons, specifically capable of selectively eliminating trypanosome parasites responsible for causing Chagas disease and African sleeping sickness (58). These DNA topoisomerase II poisons hold tremendous promise as potential therapeutics for the treatment of Chagas disease (58). Future efforts should focus on discovering poisons of DNA topoisomerases including DNA gyrase poisons.

Here we present a novel fluorescence-based, T5 exonuclease-amplified DNA cleavage assay for discovering bacterial DNA gyrase poisons. This assay can be developed into an HTS assay to discover new DNA gyrase poisons, as it allows for the screening of compound libraries comprising thousands or even millions of compounds. Additionally, similar assays could be developed to identify poisons targeting other DNA topoisomerases.

## Materials and Methods

### Proteins, plasmids, and other reagents

*E. coli* DNA gyrase, T5 exonuclease, and His-tagged human DNA topoisomerase IIα C-terminal deletion mutant (hTopo2α-ΔCTD) were purified as described previously (59,60). Plasmid pBR322 and lambda DNA HindIII digest were purchased from New England Biolabs, Inc (Ipswich, MA). Relaxed pBR322 was prepared using *variola* DNA topoisomerase I purified in our lab (61). Ciprofloxacin, etoposide, sodium dodecyl sulfate (SDS), and sarkosyl or sodium lauroyl sarcosinate (SLS) were purchased from Sigma-Aldrich, Inc. SYBR™ Green I was bought from ThermoFisher Scientific, Inc. NSC compounds were obtained from NCI DTP program (https://dtp.cancer.gov). A 50-compound library was described previously (52).

### DNA gyrase and human DNA topoisomerase II*α* mediated DNA cleavage assay

250 ng of relaxed plasmid pBR322 and 20 nM of *E. coli* DNA gyrase or human DNA topoisomerase IIα were mixed and incubated in 1× gyrase-mediated DNA cleavage buffer (20 mM Tris-HCl pH 8, 50 mM KAc, 10 mM MgCl_2_, 2 mM DTT, 1 mM ATP, 0.1 mg/mL BSA) at 37 ºC for 15 minutes in the presence of an inhibitor or compound. After the incubation, 0.2% SDS and 0.1 mg/ml proteinase K were added to the reaction mixtures to trap the topoisomerase-inhibitor-DNA complex and digest the topoisomerase, respectively, by incubating for an additional 30 min at 37 ºC. DNA samples were analyzed in 1% agarose gel containing 0.5 µg/mL ethidium bromide in 1×TAE buffer and photographed under UV light. Ciprofloxacin was used as a positive control for *E. coli* DNA gyrase-mediated DNA cleavage assays. Etoposide was used as a positive control for human DNA topoisomerase IIα-mediated DNA cleavage assays.

### A fluorescence-based, T5 exonuclease-amplified DNA cleavage assay to identify poisons for *E. coli* DNA gyrase

The fluorescence-based, gyrase-amplified DNA cleavage assays were performed in 20 µL of 1× gyrase-mediated DNA cleavage buffer (20 mM Tris-HCl pH 8, 50 mM KAc, 10 mM MgCl_2_, 2 mM DTT, 1 mM ATP, 0.1 mg/mL BSA) containing 200 ng of relaxed pBR322 and 0.05 mg/mL (136.5 nM) of *E. coli* DNA gyrase in the absence or presence of a potential gyrase poison. For positive control experiments, 50 µM of ciprofloxacin was used. For negative control experiments, 1% DMSO was used. After 60 min of incubation at 37 °C, 0.125% of sarkasyl (final concentration) was added to trap the drug-gyrase-DNA complexes. 0.5 mg/mL of proteinase K was then added to the reaction mixtures to digest DNA gyrase at 50 °C for 30 min. After the DNA samples were diluted 100-fold using 1×NEB buffer 4 (20 mM Tris-Acetate, pH 7.9, 50 mM KAc, 10 mM Mg(AC)_2_, and 1 mM DTT), 30 µL of each DNA sample was transfer to a well of a 384-well plate. 200 nM of T5 exonuclease was used to digest DNA fragment for 2 hours at 37 °C. 1×SYBR Green was added to all DNA samples. Fluorescence intensity was measured using a Biotek microplate reader with excitation wavelength of 497 nm and emission wavelength of 525 nm.

Z-factor (Z’) was determined using 96 wells of a 384-well plate where 48 wells are for positive controls in the presence of 50 µM of ciprofloxacin and the rest 48 wells for negative controls in the absence of ciprofloxacin. Z’ was calculated by the following equation:

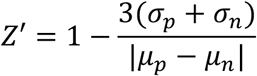

where σ_p_, σ_n_, µ_p_ and µ_n_ represent the sample means and standard deviations for positive (p) and negative (n) controls, respectively.

## Results and Discussion

Fig. 1 shows the principle and experimental strategy for a fluorescence-based, T5 exonuclease-amplified DNA cleavage assay designed to discover bacterial DNA gyrase poisons. We hypothesize that multiple DNA gyrase molecules simultaneously bind to a plasmid DNA molecule to form a gyrase-plasmid complex, which leads to the formation of multiple gyrase-DNA cleavage complexes on the same plasmid DNA molecule (Fig. 1A). In the presence of a DNA gyrase poison, such as ciprofloxacin, these gyrase-DNA cleavage complexes can be trapped by SDS or sarkosyl (sodium lauroyl sarcosinate, SLS). Following gyrase digestion by proteinase K, fragmented linear DNA molecules are generated. T5 exonuclease, which selectively digests linear and nicked DNA while leaving supercoiled and relaxed plasmid DNA intact (52), can be employed to completely digest the fragmented linear DNA molecules and “amplify” the difference of the remaining DNA molecules in the DNA cleavages assays between the presence and absence of a poisoning gyrase inhibitor (Fig. 1B). The reduction in the plasmid can be detected using SYBR green staining dye or quantitative polymerase chain reaction (qPCR). As this is a fluorescence-based assay, it can be configured into a miniaturized, automated HTS assay capable of identifying DNA gyrase poisons by screening compound libraries containing thousands or millions of compounds. Importantly, this assay not only detects gyrase poisons resulting in double-stranded DNA breaks but also identifies those causing single-stranded DNA breaks, such as NTBIs (e.g., gepotidacin) (45,46).

**Figure 1.**
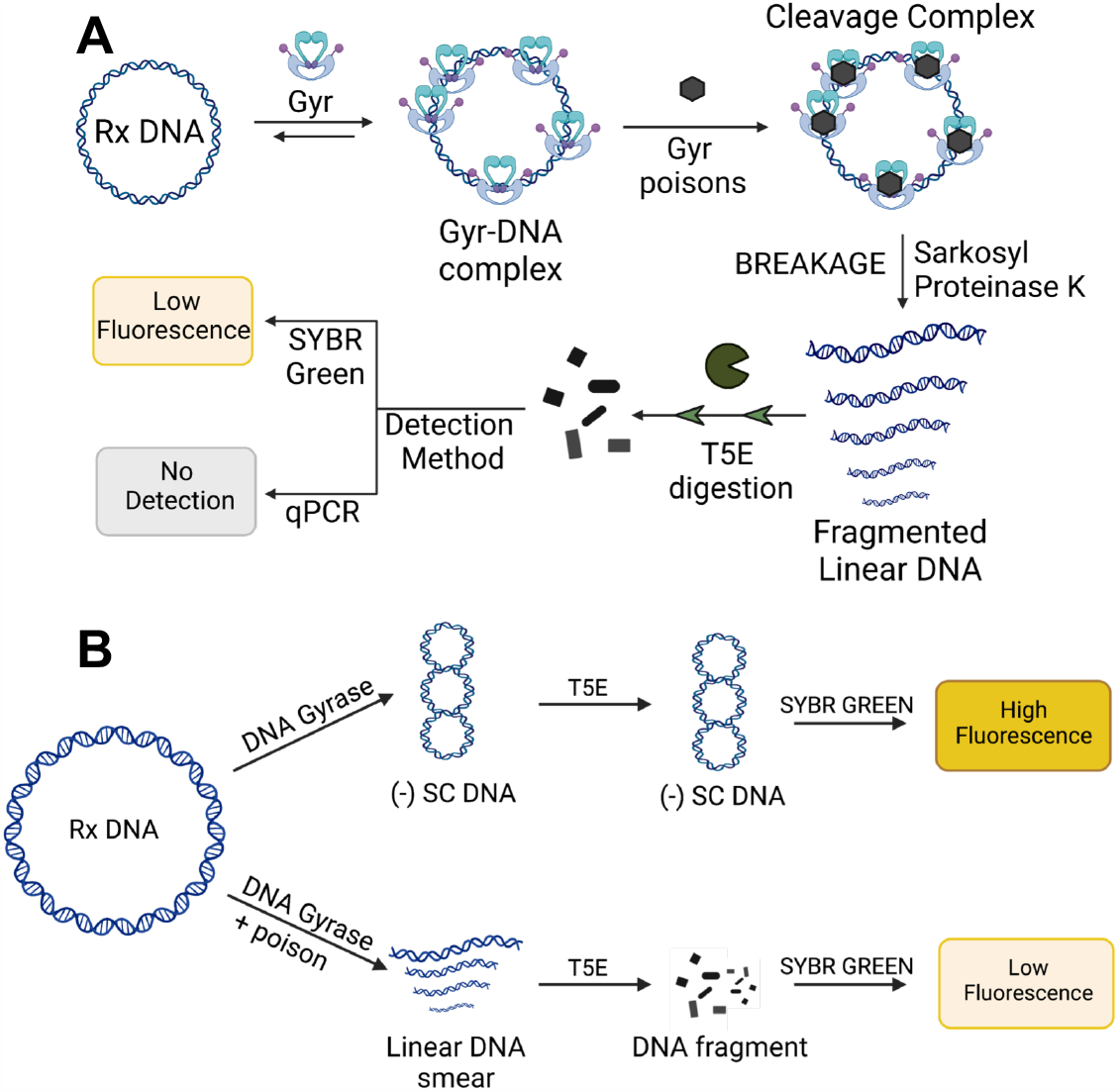
A novel fluorescence-based, T5 exonuclease-amplified DNA cleavage assay for the discovery of bacterial DNA gyrase poisons. **(A)** The principle. **(B)** The experimental strategy. This novel unique assay can be developed into automatic and miniature high throughput screening (HTS) assay to discover/identify bacterial DNA gyrase poisons by screening compound libraries that contain thousands or millions of drug-like compounds.

This fluorescence-based, T5 exonuclease-amplified DNA cleavage assay originated from an intriguing observation made during our agarose gel-based, bacterial gyrase-mediated DNA cleavage experiments. In the presence of 50 µM of ciprofloxacin and a high concentration of *E. coli* DNA gyrase, e.g., 150 nM of *E. coli* DNA gyrase, we observed the conversion of all plasmid pBR322 molecules into small DNA fragments (Fig. 2A). The fragmentation of plasmid DNA pBR322 was contingent upon the presence of ciprofloxacin (compare lane 1 to lanes 2-6 of Fig. 2A) and correlated with the concentration of ciprofloxacin (Fig. 2B). We conducted a DNA gyrase titration experiment for the gyrase-mediated DNA cleavage assays and noted a progressive increase in the production of linear plasmid DNA products with escalating concentrations of *E. coli* DNA gyrase, all in the presence of 50 µM of ciprofloxacin (Fig. 2C). Remarkably, we observed DNA smears extending from the linear band and traversing the supercoiled band when employing 26.8 and 53.6 nM of DNA gyrase (lanes 4 and 5 of Fig. 2C). Consequently, there was a significant reduction in the supercoiled DNA bands in these lanes. As anticipated, and consistent with the results presented in Fig. 2A, when 150 nM of DNA gyrase was present, the supercoiled plasmid DNA template disappeared entirely (lanes 8 and 9 of Fig. 2C). These findings provide robust support for our hypothesis that the presence of two or more gyrase molecules on a plasmid molecule leads to the formation of several gyrase-DNA cleavage complexes. These gyrase-DNA cleavage complexes, stabilized by ciprofloxacin, were subsequently entrapped by SDS or sarkosyl. Following proteinase K digestion of gyrase, fragmented linear DNA molecules were generated (Fig. 1A). Furthermore, these results serve as the foundation for the proposed fluorescence-based, T5 exonuclease-amplified DNA cleavage assay (Fig. 1B).

**Figure 2.**
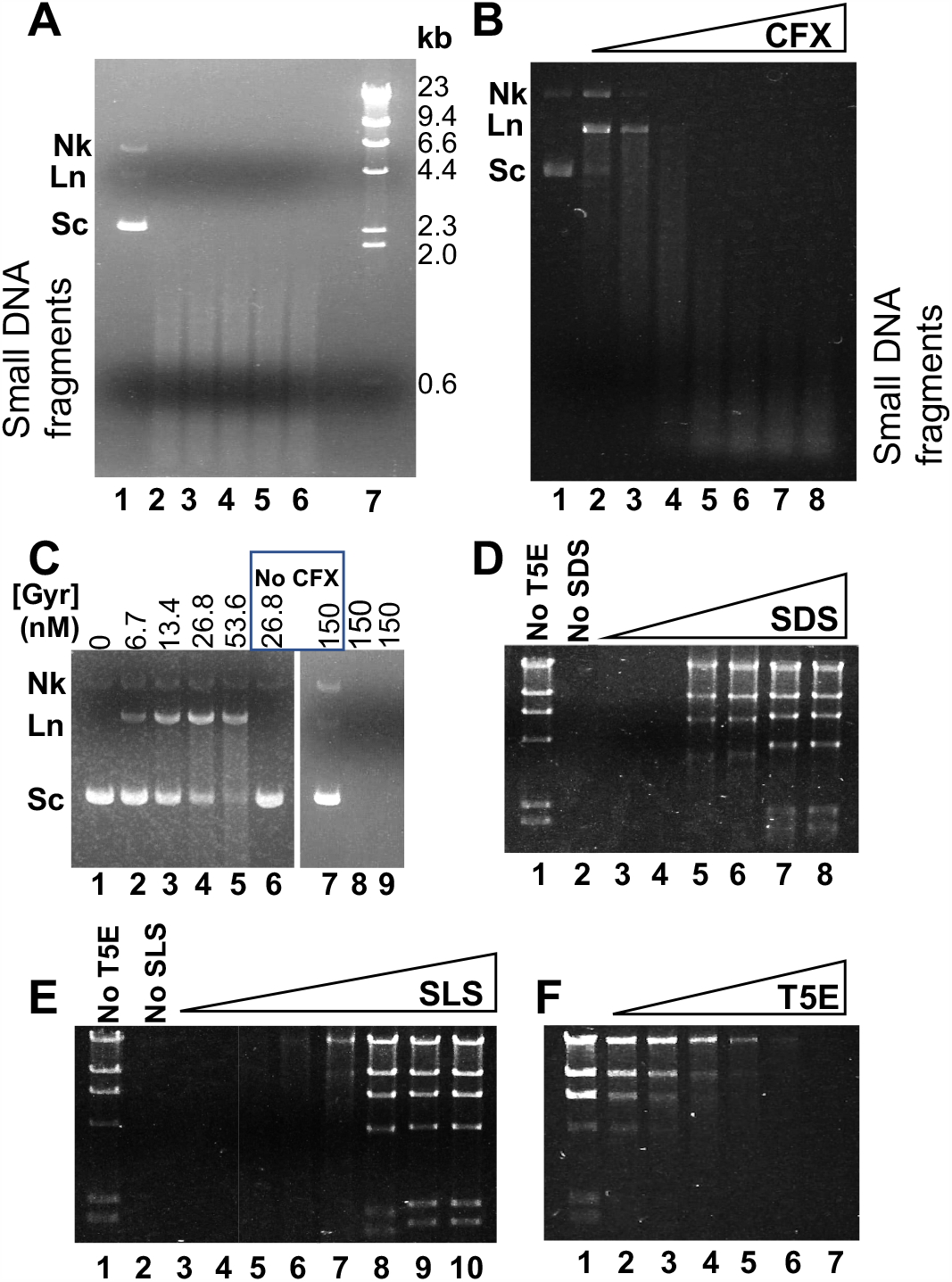
Agarose gel-based DNA gyrase cleavage assays were performed as described in Materials and Methods. **(A)** Very high concentrations of *E. coli* DNA gyrase (150 nM) cleaved plasmid pBR322 into small DNA fragments in the presence of 50 µM of ciprofloxacin in the gyrase-mediated DNA cleavage assays. Lane 1 is the DNA sample from the reaction mixture in the absence of ciprofloxacin. Lanes 2-6 contain DNA samples from the reaction mixtures containing 50 µM of ciprofloxacin. Lane 7 is Lambda DNA HindIII digest as the molecular standard. **(B)** Sufficient amount of ciprofloxacin is required for the cleavage of plasmid pBR322 into small DNA fragments. Lanes 1-8 contain 0, 0.5, 1, 2, 5, 10, 25, 50 µM of ciprofloxacin, respectively. **(C)** The appearance and disappearance of the gyrase-mediated linear band of plasmid pBR322 is dependent of the *E. coli* DNA gyrase concentration. High concentrations of *E. coli* DNA gyrase also cause disappearance of the supercoiled plasmid in the gyrase-mediated DNA cleavage assays. **(D)** SDS potently inhibited T5 exonuclease (T5E) activity. 500 ng of λ DNA HindIII digest was digested using 50 nM of T5E in 20 µL of 1× gyrase-mediated DNA cleavage buffer (20 mM Tris-HCl pH 8, 50 mM KAc, 10 mM MgCl_2_, 2 mM DTT, 1 mM ATP, 0.1 mg/mL BSA) for 30 min in the presence of different concentrations of SDS. Lanes 2-8 contained 0.000625, 0.00125, 0.0025, 0.005, 0.0125, and 0.025%, respectively. **(E)** Effects of sarkosyl on T5 exonuclease. 500 ng of λ DNA HindIII digest was digested using 50 nM of T5E in 20 µL of 1× gyrase-mediated DNA cleavage buffer for 30 min in the presence of different concentrations of sarkosyl. Lanes 3 to 10 contained 0.005, 0.0125, 0.015, 0.02, 0.025, 0.05, 0.125, and 0.25%, respectively. **(F)** 50 nM of T5 exonuclease completely digested 500 ng of λ DNA HindIII digest in 1× gyrase-mediated DNA cleavage buffer in the presence of 0.00125%. Lanes 1-7 contained 0, 5, 10, 20, 30, 40, and 50 nM of T5 exonuclease, respectively.

In a typical gyrase-mediated DNA cleavage assay, SDS or sarkosyl is used to trap the drug-gyrase-DNA complexes (50,62). Our results showed that low concentrations of SDS completely inhibited T5 exonuclease activity (Fig. 2D). In contrast, relatively high concentrations of sarkosyl (e.g., 0.015%) did not significantly affect T5 exonuclease’s activity (Fig. 2E). Therefore, we chose to use sarkosyl to trap the drug-gyrase-DNA complexes in subsequent assays. Indeed, in the presence of sarkosyl, 50 nM of T5 exonuclease successfully digested 250 ng of lambda DNA HindIII digest within 30 minutes (Fig. 2F).

A series of experiments were conducted to determine the optimal conditions for the fluorescence-based, T5 exonuclease-amplified DNA cleavage assay for *E. coli* DNA gyrase (Fig. 3). For example, in the presence of ciprofloxacin, the fluorescence intensity of the DNA samples progressively decreased as the concentration of *E. coli* DNA gyrase was increased (Fig. 3A). It reached a plateau when 100 nM of *E. coli* DNA gyrase was used (Fig. 3A). In contrast, the concentration of *E. coli* DNA gyrase did not significantly change the fluorescence intensity for the DNA samples in the absence of ciprofloxacin (Fig. 3A). Additionally, the fluorescence intensity of the DNA samples depended on the concentration of ciprofloxacin (Fig. 3B). T5 exonuclease greatly “amplified” the fluorescence difference/signal of the DNA samples between the presence and absence of ciprofloxacin (Fig. 3C and D). Diluting the DNA samples greatly reduced the effects of sarkosyl on the digestion of DNA samples by T5 exonuclease (Fig. 3E). 1× and 2× SYBR Green provided the best results for the HTS assays (Fig. 3F). These experiments established that the optimal conditions for the HTS assay included the use of 136.5 nM of *E. coli* DNA gyrase, 50 µM of ciprofloxacin (for positive controls), 200 nM of T5 exonuclease, a 100-fold dilution of DNA samples, and 1× SYBR Green. The assay demonstrated tolerance to up to 2% DMSO without any significant change in signal. In a titration experiment in which different concentrations of ciprofloxacin were added to the assays, it became evident that ciprofloxacin progressively cleaved DNA, with estimated EC50 values of 1.48 µM (Fig. 3B). This EC50 value is consistent with previously published results (56).

**Figure 3.**
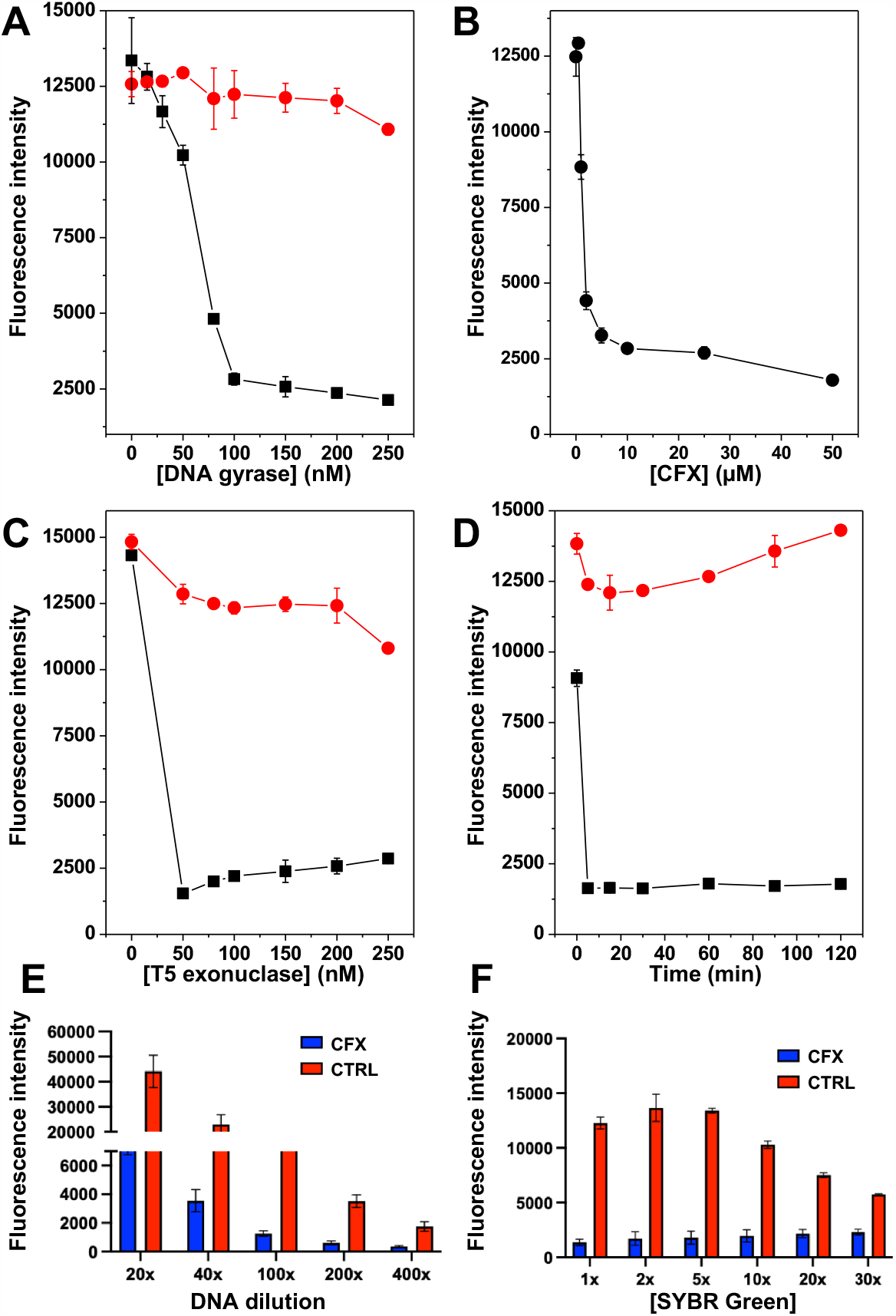
Determining the optimal conditions for the fluorescence-based, T5 exonuclease-amplified DNA cleavage assay for *E. coli* DNA gyrase. The fluorescence-based, T5 exonuclease-amplified DNA cleavage assay for *E. coli* DNA gyrase was described in Materials and Methods. The fluorescence intensity was measured with λ em = 525 nm and λ ex = 497 nm using a plate reader. **(A-D)** Red and black dots-lines represent assays in the absence and presence of ciprofloxacin (CFX). **(A)** *E. coli* gyrase titration experiments. **(B)** Ciprofloxacin titration experiment. **(C)** T5 exonuclease titration experiment. **(D)** Time courses for T5 exonuclease digestion. 200 nM of T5 exonuclease was used in each assay. **(E)** Dilution effect of DNA samples before T5 exonuclease digestion. **(F)** SYBR Green titration experiment.

To validate our assay, we conducted a preliminary experiment with ten compounds, comprising five fluoroquinolones (FQs) and five other antibiotics (novobiocin, ampicillin, polymyxin B, sulfanilamide, and kanamycin). The results are shown in Fig. 4A and B. Notably, the fluorescence intensity of the DNA samples in the presence of FQs was significantly low, whereas the fluorescence intensity of the DNA samples in the presence of other antibiotics, including novobiocin, was notably high. Interestingly, novobiocin, known as a DNA gyrase catalytic inhibitor targeting the gyrase ATPase (61,63-65), did not induce a decrease in fluorescence intensity. These findings demonstrate the feasibility of our assay, which is tailored for the identification of DNA gyrase poisons.

**Figure 4.**
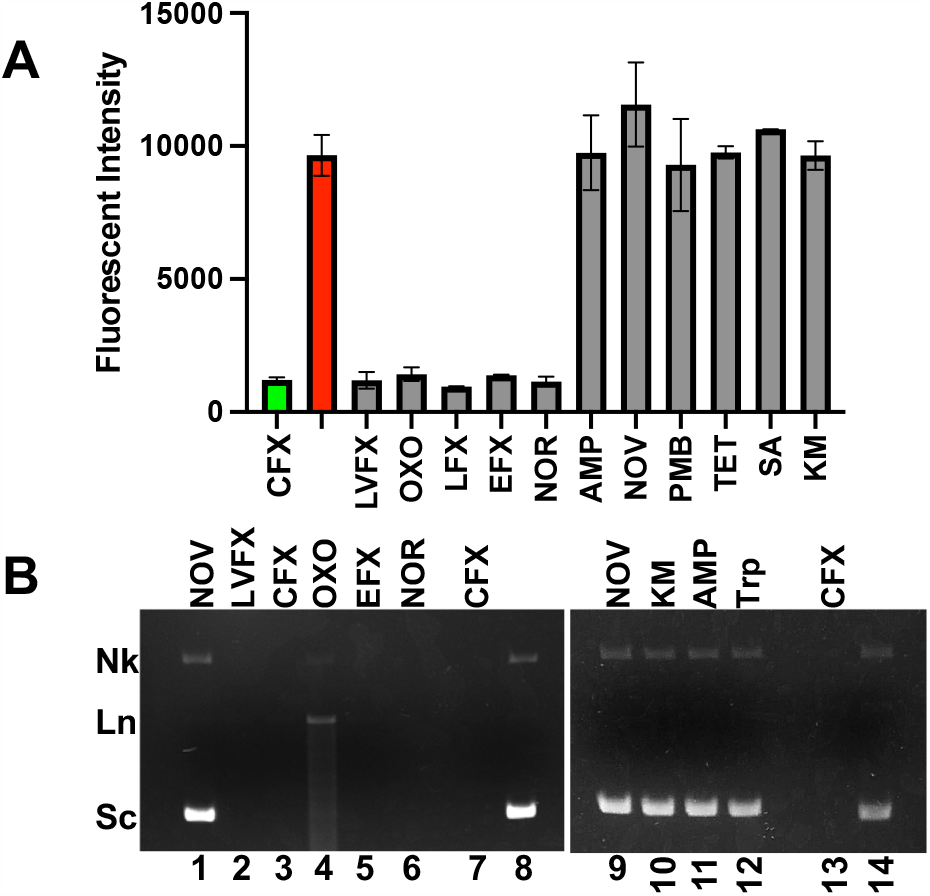
The fluorescence-based, T5 exonuclease-amplified DNA gyrase cleavage assay for 10 different antibiotics, 5 fluoroquinolones and 5 other antibiotics were performed in 20 µL of 1× gyrase-mediated DNA cleavage buffer containing 200 ng of Rx pBR322 and 0.05 mg/mL (136.5 nM) of *E. coli* DNA gyrase in the absence or presence of an antibiotics. For control experiments, 50 µM of ciprofloxacin was used. After 60 min of incubation at 37 °C, 0.125% of sarkasyl was added to trap the drug-gyrase-DNA complexes. 0.5 mg/mL of proteinase K was then added to the reaction mixtures to digest DNA gyrase at 50 °C for 30 min. After the DNA samples were diluted 100-fold using 1×NEB buffer 4, 200 nM of T5 exonuclease was used to digest DNA fragment for 2 hours at 37 °C. 1×SYBR Green was added to all DNA samples. **(A)** Fluorescence intensity was measured using a Biotek microplate reader with excitation wavelength of 497 nm and emission wavelength of 525 nm. **(B)** 1% agarose gels containing 0.5 µg/mL of EB in 1×TAE for DNA samples before the 100 times of dilution. Lanes 8 and 14 did not contain a compound. Abbreviations: Nov, novobiocin; CFX, ciprofloxacin; OXO, oxolinic acid; LFX, lomefloxacin; EFX, enrofloxacin; NOR, norfloxacin; AMP, ampicillin; PMB, polymyxin B sulfate; TET, tetracycline SA, sulfanilamide; KM, kanamycin.

Next, we screened a 50-compound library assembled in our lab (52) using this assay at a final compound concentration of 50 µM in duplicate and got the following statistics: *Z*′, 0.53; *S*/*B*, 7.3; and 8 hits (Fig. 5, Table 1, and Table S1). We chose 50 µM since we always used 50 µM of ciprofloxacin as the positive controls. We also wanted to test whether the fluorescence from some of these compounds in the library interfere with the assay, as previous HTS assays for DNA topoisomerase inhibitors had been significantly affected by certain compounds with strong fluorescence (50,52). Our results clearly demonstrated that fluorescence from these compounds per se did not interfere with the fluorescence signal from SYBR Green staining (Fig. 5 and Table S1).

**Table 1.** Parameters for the fluorescence-based, T5 exonuclease-amplified HTS assay for *E. coli* DNA gyrase using the 50-compound library.

|  |  |
| --- | --- |
| # of compounds | 50 |
| Tested concentration | 50 μM |
| Z' value | 0.53 |
| <sup>a</sup>S/B | 7.3 |
| # of compounds<br>with inhibition>50% | 8 |

<sup>a</sup>S/B represents signal versus background ratio.

**Figure 5.**
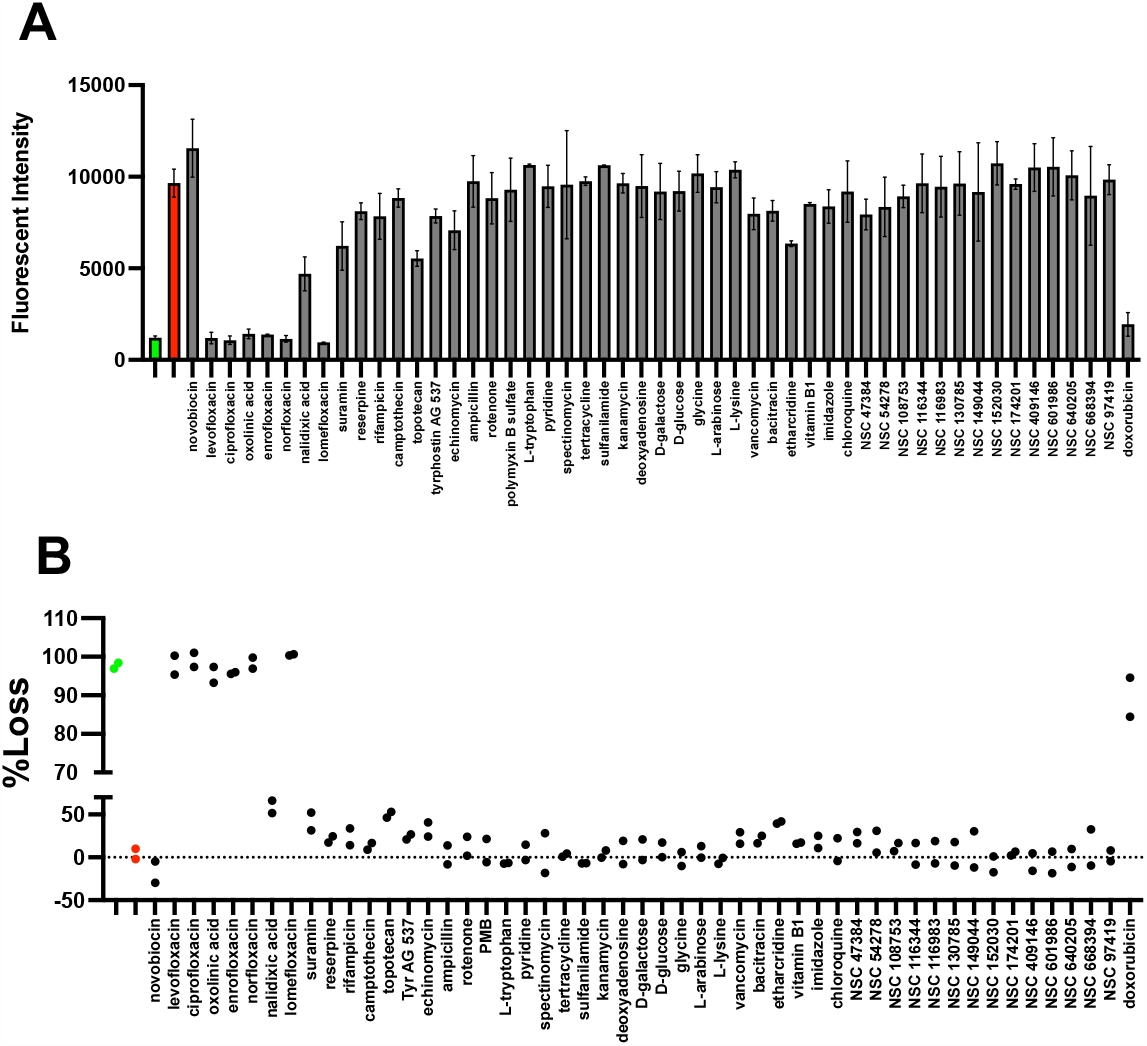
High throughput screening (HTS) pilot screen of the 50-compound library for *E. coli* DNA gyrase poisons in duplicate. A final compound concentration of 50 µM was used. DMSO (1%) (red) and ciprofloxacin (50 µM) (green) are used as negative and positive controls, respectively. **(A)** Fluorescence raw data. **(B)** Decrease in DNA signal is calculated by using equation 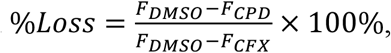, where F_DMSO_, F_CPD_, and F_CFX_ are fluorescence intensity of DNA samples from DNA cleavage assays containing 1% DMSO (as negative controls), a compound (CPD), and ciprofloxacin (CFX), respectively. A loss of ≥50% of DNA fluorescence signal against *E. coli* DNA gyrase is used as the cutoff value for gyrase poisons, which results in 8 hits.

Among the eight hits, seven were quinolones (including levofloxacin, ciprofloxacin, oxolinic acid, enrofloxacin, norfloxacin, lomefloxacin, and nalidixic acid), along with doxorubicin. Quinolones are well-known as potent gyrase poisons (20,21). It was unsurprising to rediscover them as such in this HTS assay. Interestingly, even though nalidixic acid exhibited an IC50 value significantly higher than 50 µM (66), the fluorescence intensity of the DNA samples in the presence of nalidixic acid was considerably lower compared to the negative controls. In essence, this HTS assay is capable of identifying gyrase poisons, even if they possess a high IC50 against *E. coli* DNA gyrase.

The only non-quinolone hit is doxorubicin, a DNA intercalator tightly binding to DNA (67). Agarose gel-based DNA cleavage assays showed that doxorubicin did not cause gyrase-mediated DNA cleavage, thus excluding it as a gyrase poison (Fig. S1). In other words, doxorubicin represented a false positive. One plausible explanation is that the high affinity of doxorubicin for DNA resulted in the quenching of SYBR Green’s fluorescence. To further explore this, we repeated the fluorescence-based DNA cleavage assay using a concentration of 20 µM, once again observing the quenching of SYBR Green fluorescence (Fig. S1A). Subsequently, we conducted a doxorubicin titration experiment, titrating different concentrations of doxorubicin into 200 ng of Rx pBR322 within a 1× gyrase DNA cleavage buffer. After diluting the reaction mixtures 100-fold, we added 1× SYBR Green. The results presented in Fig. S1C clearly confirmed that doxorubicin indeed caused the quenching of SYBR Green fluorescence.

In this pilot screen, we observed that several compounds, including ethacridine, echinomycin, topotecan, and suramin, caused notably low fluorescence signals in the DNA samples from the DNA cleavage assays conducted at 50 µM. However, when the final compound concentration was reduced to 20 µM, these compounds no longer resulted in a reduction in the fluorescence intensity of SYBR Green. This observation suggests that they do not function as gyrase poisons (Fig S1A). Further validation through agarose gel-based DNA cleavage assays corroborated these findings, confirming that indeed, these compounds do not possess gyrase poisoning properties (Fig S1B).

We also explored the potential of developing a similar DNA cleavage assay for human DNA topoisomerase IIα, given that several anticancer drugs, such as etoposide and doxorubicin, serve as poisons targeting this specific human DNA topoisomerase (8). Fig. S2 shows our results. A titration experiment of human DNA topoisomerase IIα showed a progressive increase in the linear and nicked plasmid DNA products when using increasing concentrations of topoisomerase IIα in the presence of 50 µM of etoposide (Fig. S2A). However, we encountered challenges in converting all plasmid DNA into the linear or nicked form (Fig. S2). One possibility is that the IC50 of etoposide against human DNA topoisomerase IIα is higher than 50 µM (68). In such cases, not all plasmid DNA molecules could form topoisomerase IIα-DNA cleavage complexes and subsequently transform into linear or nicked forms, even after SDS or sarkosyl had entrapped the cleavage complexes and proteinase K had digested topoisomerase IIα.

Furthermore, we observed that in the absence of phenol extraction, certain DNA-protein complexes were unable to migrate into the agarose gel and remained trapped in the wells (Fig. S2B). Nevertheless, we did detect a significant difference in fluorescence intensity between the DNA samples in the presence and absence of etoposide (Fig. S2C). These results underscore the feasibility of establishing a similar fluorescence-based DNA cleavage assay for human DNA topoisomerase IIα. However, we opted not to pursue this avenue further, given that the primary focus of this study was to establish a fluorescence-based, T5 exonuclease-amplified DNA cleavage assay for bacterial DNA gyrase. Moreover, the optimization of the fluorescence-based DNA cleavage assay would require a substantial quantity of human DNA topoisomerase IIα, which is not readily available in our laboratory.

In summary, we have established a novel fluorescence-based DNA cleavage assay to discover bacterial DNA gyrase poisons, based on a recent observation showing that multiple DNA gyrase molecules simultaneously bind to a plasmid DNA molecule to form multiple gyrase-DNA cleavage complexes on the same plasmid DNA molecule. These gyrase-DNA cleavage complexes, greatly stabilized by a DNA gyrase poison, can be trapped by SDS or sarkosyl. Digestion of DNA gyrase by proteinase K result in the production of small DNA fragments. T5 exonuclease, which selectively digests linear and nicked DNA, can be used to completely digest the fragmented linear DNA molecules and, therefore, “amplify” the fluorescence signal of the DNA cleavage products after SYBR Green staining. We validated this fluorescence-based DNA cleavage HTS assay using a 50-coupound library. It has a Z prime value more than 0.5 and ready to screen different compound libraries. Similar fluorescence-based DNA cleavage assays could also be established to identify poisons against other DNA topoisomerases, such as human topoisomerase IIα.

## Supplementary Information

Supplementary Information is available at *Nucleic Acids Research* Online.

## Acknowledgments

We thank Dr. Yuk-Ching Tse-Dinh for critically reading the manuscript before submission. This work was supported by National Institutes of Health grants 1R21AI125973 and 1R21AI178134 (to F.L.).

## Author contribution

F.L. designed research; M.D. and T.C. performed research; F.L. and M.D. analyzed data; F.L. wrote the paper.

---

**Figure S1.**
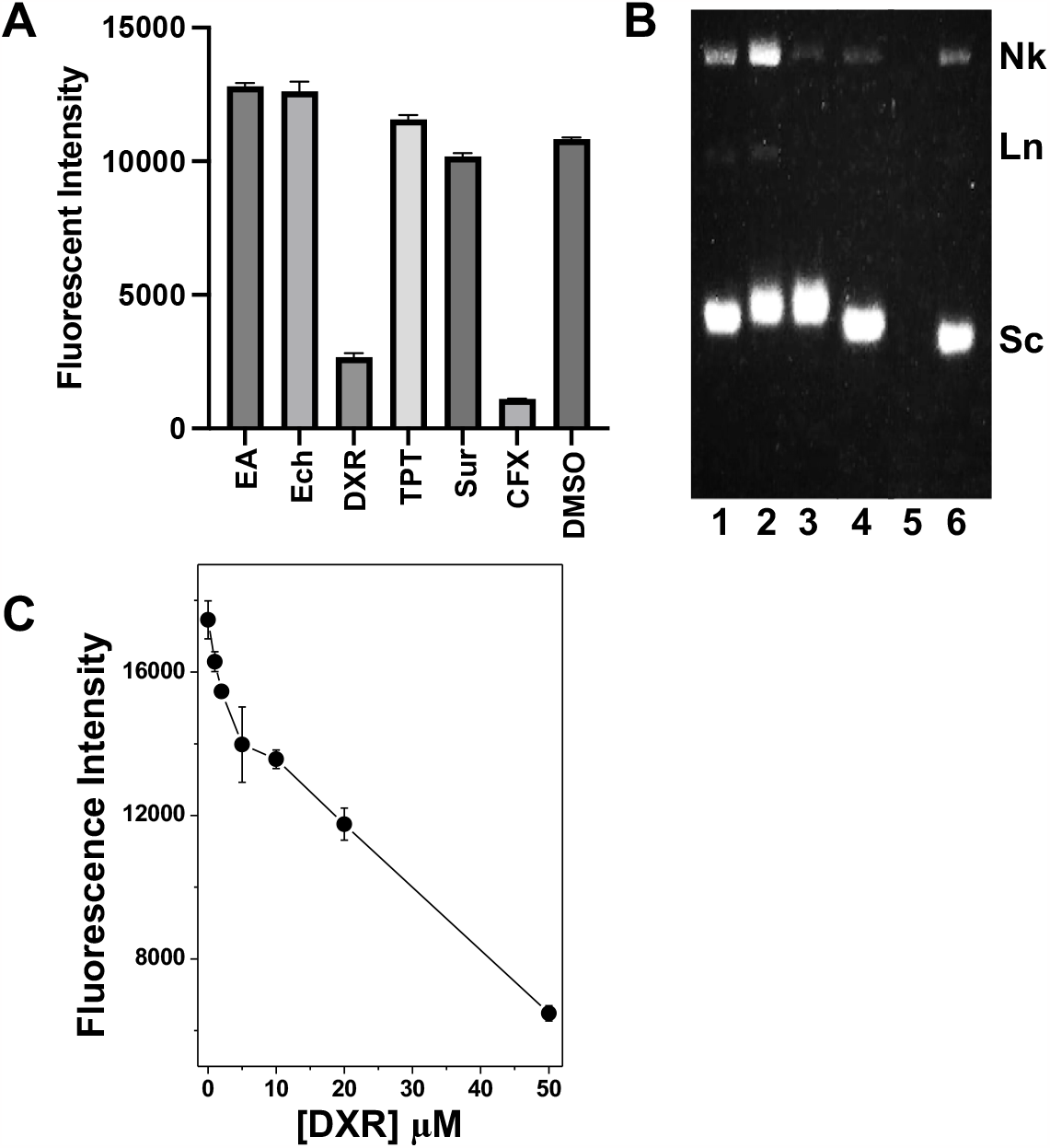
The fluorescence-based, T5 exonuclease-amplified DNA gyrase cleavage assays for ethacridine (EA), echinomycin (Ech), doxorubicin (DXR), topotecan (TPT), and suramin (Sur) at 20 µM were formed as described in Materials and Methods. **(A)** Fluorescence intensity was measured using a Biotek microplate reader with λex of 497 nm and λem of 525 nm. **(B)** 1% agarose gels containing 0.5 µg/mL of EB in 1×TAE for DNA samples before dilution. Lanes 1-5 are DNA samples from assays containing ethacridine, echinomycin, doxorubicin, suramin, and ciprofloxacin, respectively. Lane 6 is the DNA sample from an assay containing 1% DMSO as a negative control. **(C)** Doxorubicin (DXR) efficiently quenched fluorescence of SYBR Green. DXR titration assays were performed in 20 µL of 1×DNA cleavage buffer containing 200 ng of Rx pBR322 and different concentrations of DXR. The DNA samples were diluted 100 times using 1×NEB buffer 4. 1×SYBR Green was added to all DNA samples. Fluorescence intensity was measured with λex of 497 nm and λem of 525 nm.

**Figure S2.**
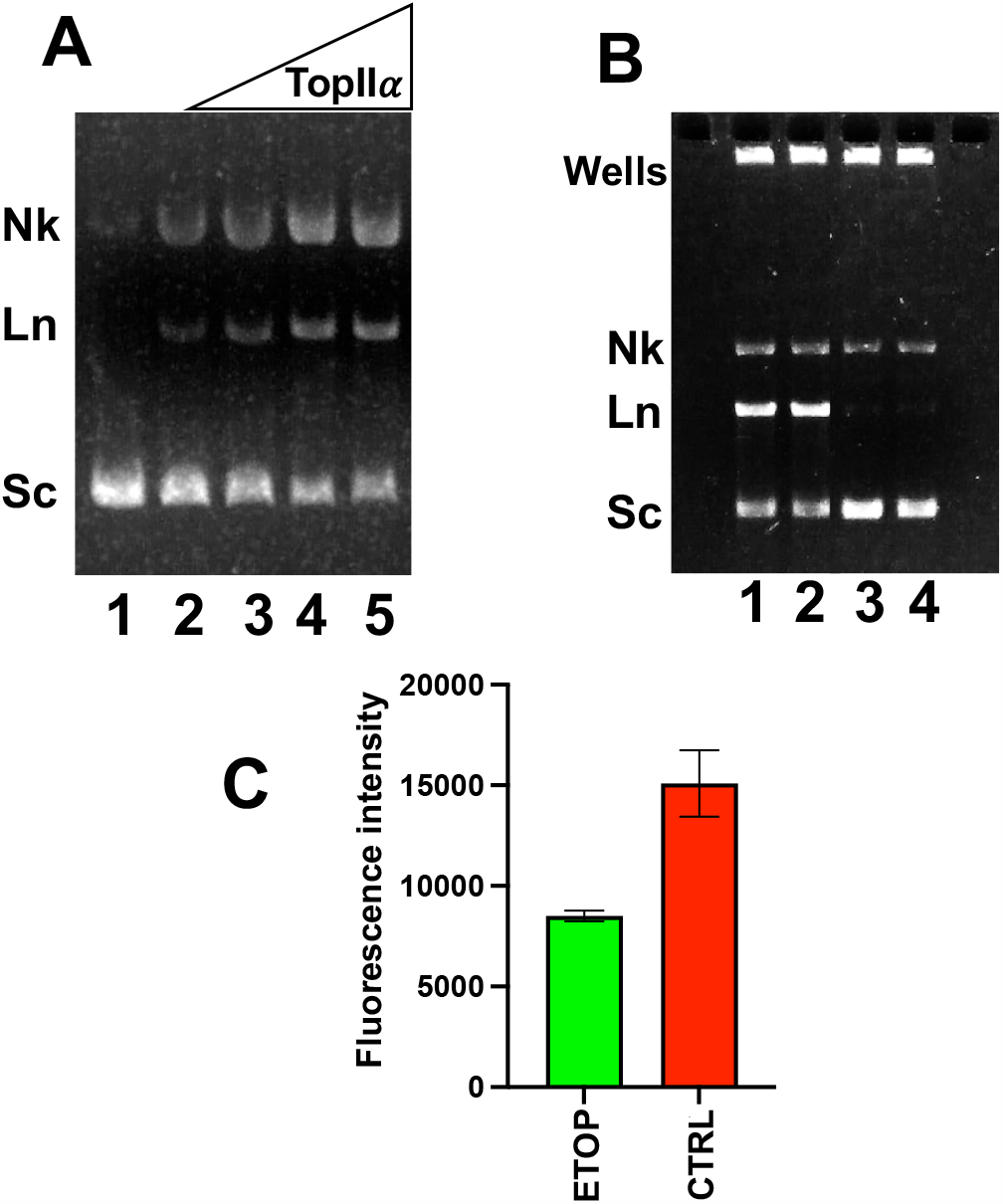
Human DNA topoisomerase II α (TopoIIα) mediated DNA cleavage assays, similar to the gyrase-mediated DNA cleavage assay, were performed as described in Materials and Methods. **(A)** and **(B)** Agarose gel-based DNA gyrase cleavage assays. **(A)** In the presence of 50 µM etoposide, human TopoIIα mediated DNA cleavage products (the linear DNA and nicked DNA) are proportional to the amount of human TopoIIα used in the assay. Phenol extraction was used to purify DNA samples before the gel electrophoresis. Lanes1-6 contain 0, 0.015, 0.03, 0.06, and 0.12 mg/mL of human TopoIIα, respectively. **(B)** Without phenol extraction, some DNA-protein complexes could not migrate from the wells into the gel. Lanes 1-4 are DNA samples from the assays in the presence (lanes 1 and 2) or absence (lanes 3 and 4) of 50 µM etoposide. 0.12 mg/mL of human TopoIIα was used for each assay. **(C)** Fluorescence intensity of the DNA samples in **(B)** after T5 exonuclease digestion and SYBR Green staining. Fluorescence intensity was measured using a Biotek microplate reader with excitation wavelength of 497 nm and emission wavelength of 525 nm.

